# A unique bacterial family strikes again!

**DOI:** 10.1101/2024.10.01.615302

**Authors:** Gyanesh Shukla, Gaurav Sharma

## Abstract

In a recent study, Rolf Müller and colleagues identified a novel myxobacterial family Pendulisporaceae, encompassing four new strains with biosynthetic gene clusters (BGCs) largely distinct from known families. These strains, sourced from 1980s soil samples, display unique genomic and chemical features, suggesting significant biotechnological and pharmaceutical potential. This study underscores the value of exploring underrepresented microbial taxa for novel natural products, highlighting the potential of the family Pendulisporaceae as a source of new antimicrobial and therapeutic agents.

## Introduction

The golden age of antibiotics commenced in the 1940s with the discovery of streptomycin, a secondary metabolite produced from *Streptomyces griseus*, a soil-borne class Actinomycetes member of the phylum Actinomycetota. This era, which was marked by rapid advancements in antibiotic discoveries, concluded two decades later, around the 1960s ^1,2^. In those twenty years, researchers globally endeavored to isolate novel microorganisms from diverse ecological niches, leading to the identification of diverse antibiotic classes. However, post-1960s, the majority of screened organisms, although taxonomically diverse at species, genus, family, and class level, yielded compounds, similar to those previously identified ^3^. These organisms were able to grow in the lab which allowed researchers to identify their novel secondary metabolites. Despite extensive global efforts, only about 1% of the Earth’s microbial diversity has been cultivated and studied, with the remaining 99% comprising viable but non-culturable bacteria and yet-to-be-cultivated species ^2,4^. The ongoing metagenomic studies from any ecological sample also corroborate this as the large plethora of reads coming from unculturable organisms remains unclassified in those studies. Over the past few decades, culture-independent molecular techniques, including denaturing and temperature gradient gel electrophoresis, terminal restriction fragment length polymorphism, 16S rDNA clone library preparation, fatty acid methyl esters analysis, and next-generation sequencing technologies, have proven effective in elucidating the genetic diversity and community structures of microorganisms, including unculturable bacteria. Owing to improved sequencing quality and throughput along with a massive decrease in costs, genome mining approaches can help in prioritizing studying microbial communities for metabolite screening and further facilitating pathways for synthetic biology investigations. Genome-mining approaches can prioritize microbial communities for screening and facilitate pathways for synthetic biology investigations ^5^. Nonetheless, the extraction of natural compounds from these unculturable microorganisms remains a significant challenge.

Myxobacteria belong to a distinctive phylum Myxococcota ^6^, notable for their transition from unicellular to multicellular lifestyles. These organisms exhibit complex social behaviors such as cooperative motility via social and adventurous mechanisms, multicellular fruiting body formation, mass predation, kin selection, and biofilm formation. This phylum is curiously known for encompassing an extremely large genome that ranges from 9-16 Mb. The majority of these organisms are soil-dwelling in nature and therefore, they synthesize a diverse array of secondary metabolites. These biosynthetic gene clusters (BGC) encoded secondary metabolites secreted by the whole swarm potentially facilitating co-operative mass predation. These organisms secrete a large plethora of polyketide synthase (PKS), non-ribosomal peptide synthetase (NRPS), hybrid PKS-NRPS, along with several other terpenes, ribosomally synthesized and post-translationally modified peptide (RiPP), alkaloids, etc ^7,8^.

Several well-known myxobacterial metabolites have been identified in the last two decades, including **epothilone** (a potent anticancer agent stabilizing microtubules similar to Taxol), **myxalamide** (cytotoxic agent disrupting cellular respiration by interfering with the electron transport chain), **sorangicin** (an antibiotic effective against multi-drug resistant bacteria by inhibiting bacterial RNA polymerase), **sandacrabin** (potent antibiotic against Gram-positive bacteria by blocking bacterial cell wall synthesis or membrane integrity), **soraphen** (an antifungal/anticancer agent inhibiting acetyl-CoA carboxylase), **disorazol** (potential anticancer properties by disrupting microtubule formation), **myxovirescin** (potent antibiotic agent against Gram-positive bacteria by inhibiting cell wall synthesis), **chivosazol** (an antifungal disrupting the actin cytoskeleton), and **haliangicin** (antifungal and cytotoxic activities) ^7^. Many of these compounds are currently under investigation for potential therapeutic applications.

Phylum Myxococcota encompasses two classes, four orders, and nine families comprising 31 genera and >200 species. Genome analysis of several myxobacterial genomes has revealed the abundance of these BGCs ^9^; however, many of those compounds cannot be extracted in normal lab conditions owing to their uncultivability, lesser BGC expression, and cryptic nature ^2^. Therefore, it is highly relevant to identify and cultivate novel myxobacteria which can be processed through an integrated genome sequencing and chemical screening (metabolomics) pipeline to overall increase the discovery of unique and potential antibiotics.

Our *in-silico* analysis using BGC regions data available in open-access from the antiSMASH database ^10^ revealed that myxobacteria encode around 28 BGC regions on a median scale for 59 myxobacterial organisms, which is almost 3.5 times as compared to the well-known secondary metabolite producer phyla, i.e., Actinomycetota, Acidobacteriota, Planctomycetota, and Cyanobacteriota **(Figure 1A and Table S1)**. At the genus level, fourteen myxobacterial genera rank among the top organisms encoding >30 BGC regions **(Table S2)**. We observed that amongst these highest BGC-encoding organisms, phylum Actinomycetota have the most representatives with genus *Streptomyces* (772 organisms), *Nocardia* (60 organisms), *Amycolatopsis* (48 organisms) being the most explored ones. Besides, phylum Actinomycetota and Myxococcota, only two additional members, i.e., *Sulfidibacter* and *Moorena* from Acidobacteriota and Cyanobacteriota, respectively, are among these >30 BGC-encoding organisms. As PKS-NRPS and their hybrids are majorly involved in diverse antibiotics production, we further investigated their combined distribution which revealed that myxobacteria are a prolific source of these PKS-NRP-hybrids as the list of top organisms encoding >14 BGC (PKS-NRP-hybrids) regions is predominated by fourteen myxobacterial genera having 40 strains **(Figure 1B and Table S3)**.

**Figure-1:**
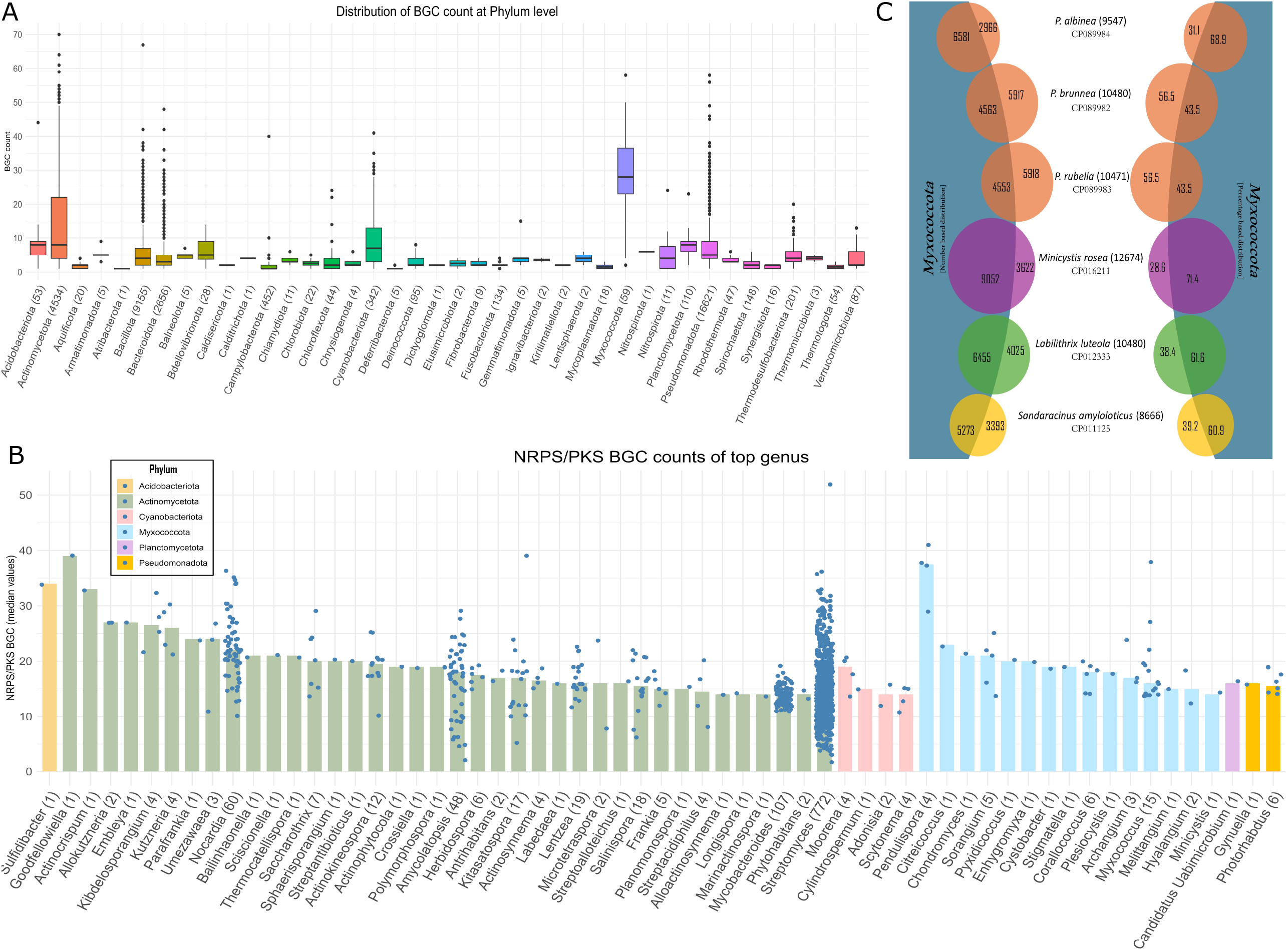
**(A) The distribution of BGC regions at the Phylum level**. The X-axis indicates the phylum name followed by the number of organisms analyzed within that phylum in parentheses, while the Y-axis represents the distribution of BGCs. **(B) The distribution of NRPS-PKS-hybrid regions at the Genus level:** Data from the antiSMASH database, encompassing over 34,000 organisms, was used to identify BGC regions, from which NRPS-PKS-hybrid regions were filtered. The bar plot displays genera encoding more than 14 NRPS-PKS-hybrid regions at the median level, along with their respective counts of NRPS-PKS-hybrid regions per genome. The X-axis represents the genus name followed by the number of organisms analyzed within the corresponding phylum in parentheses, while the Y-axis represents the distribution of NRPS-PKS-hybrid regions. The grouping is based on the phylum of each genus, as indicated in the upper left corner. **(C) The distribution of unique genes per organism at gene number and percentage level**: Protein-ortho analysis was performed in a binary manner using two datasets: 1) all encoded proteins for respective organisms as mentioned in the middle and 2) a combined dataset of all encoded proteins from 82 myxobacterial organisms. This analysis identified the number of unique genes absent in the 82 myxobacterial genomes. The organism name, along with its total encoded proteins and NCBI genome ID, is listed in the middle.

Overall, it can be argued that despite possessing a substantial array of BGCs, especially PKS-NRP-hybrids, myxobacterial members have been less extensively studied as compared to Actinobacteria. This is further evident by their predominance at the genome level, i.e., 3,560 genome assemblies for *Streptomyces*, 54,597 for phylum Actinomycetota, and only 4,191 for phylum Myxococcota. This number for myxobacteria looks higher, although, 3,861 out of 4,191 assemblies have not been assigned a proper genus name, and another 170 have not been categorized at species level; overall leaving only 160 assemblies to have proper classification with complete or draft nature.

In a study published in the June/July issue of Chem, Rolf Müller and colleagues identified a novel family of myxobacteria, Pendulisporaceae, comprising four new strains ^11^. These strains harbor biosynthetic gene clusters (BGCs), 90% of which are distinct from known BGC families, exhibiting unique chemistry and enzymology. Rolf Müller’s group at Helmholtz Institute for Pharmaceutical Research, Saarland, Germany, has been actively involved in diverse expeditions aimed at discovering novel myxobacterial strains and characterizing their secondary metabolites using metabolomic approaches. This research is a significant example of their persistence and dedication in identifying novel myxobacteria as these four strains were isolated from soil samples collected in the 1980s from Zambia and the Philippines by Prof. Hans Reichenbach, a well-known microbiologist and taxonomist.

Rolf Müller and colleagues classified the family Pendulisporaceae within suborder Sorangiineae, which has several unique myxobacteria such as the highest genome size bacterial genome (16 Mb having *Minicystis rosea*), the well-known metabolite-producer myxobacteria (11-14.8 Mb having *Sorangium cellulosum spp*.) and the only pathogenic myxobacteria with a mere 1.8 Mb genome (*Pajaroellobacter abortibovis*). It must be noted that the closest relative of the family Pendulisporaceae, *Labilithrix luteola* DSM-27648 is also a member of another novel family, Labilitrichaceae, identified nine years ago in 2014, isolated from forest soil in Kagoshima Prefecture, Japan in 2012 ^12^ and sequenced in 2015 (<u>CP012333.1</u>; unpublished data). Similarly, *Sandaracinus amylolyticus* DSM 53668 is another close relative of these organisms, which also belongs to a unique family identified by Rolf Müller’s group in 2012 from a soil sample collected in 1995 from Lucknow, Uttar Pradesh, India ^13^, and sequenced in 2016 ^14^. *Minicystis rosea* DSM 24000 is another close relative that was isolated from a Philippine soil sample collected in 2007 by Rolf Müller’s group followed by its characterization in 2014 ^15^ and sequenced in 2017 ^16^.

The four strains discussed in this Chem article, i.e., *Pendulispora rubella* MSr11367^T^ and MSr11368, *P. albinea* MSr11954^T^, and *P. brunnea* MSr12523^T^ were classified in a novel family, i.e., Pendulisporaceae, using 16S rRNA phylogeny, DNA-DNA hybridization (DDH) analysis, and dot plots. To further showcase that this family and these strains are novel, we compared their pangenome and unique genome with 85 (82 + *L. luteola* DSM-27648, *S. amylolyticus* DSM 53668, and *M. rosea* DSM 24000) available myxobacterial genomes in a binary distribution of respective genome and rest of the group **(Figure 1C)**. This comparative study with 82 organisms revealed that >56% of the encoded genes in *P. rubella* and *P. brunnea* are unique whereas *P. albinea* possesses only 31% unique genes. Additionally, *L. luteola* DSM-27648, *S. amylolyticus* DSM 53668, and *M. rosea* DSM 24000 exhibit 38%, 39%, and 28% unique genes, compared to the 82 common myxobacterial genomes. When all four *Pendulispora* genome annotations were compared against 85 myxobacterial organisms, our analysis depicted that 33.10% of genes are still unique in *Pendulispora* spp. This pan-genome analysis highlights the family *Pendulisporaceae*’s substantial uniqueness at the gene/protein homology level as compared to the previously characterized novel organisms, thereby strengthening the findings reported by Rolf Müller and colleagues.

Rolf Müller and colleagues further focus on the novel nature of their natural products (NPs) based on genomic and bioactivity-metabolite correlation investigations. The identification of these unique NPs underscores the potential of Pendulisporaceae as a source of novel bioactive compounds with significant implications for biotechnology and pharmaceuticals.

<u>Desferrioxamine PS</u>, a hydroxamate-type siderophore, structurally resembles those from Streptomyces, potentially enhancing its binding affinity or specificity for certain metal ions, and broadening its therapeutic scope. <u>Pendulipeptides A and B</u>, isolated from *Pendulispora rubella* MSr11367^T^, are N-terminally acetylated and C-terminally reduced tetrapeptides, featuring unique modifications not commonly found in bacterial peptides. A notable aspect of these NPs is their nonribosomal peptide synthetase (NRPS) system, where the installation of an acetyl moiety at the N-terminus is mediated by the monomodular *pendA* gene, which is homologous to an Ebony NRPS in *Drosophila melanogaster* and includes an atypical C_N_ domain, entirely unprecedented in bacteria. This novel enzymatic function could expand the chemical diversity of bacterial peptides. <u>Sorangicin P</u> features an additional bicyclic system unprecedented within the sorangicin family, known for its potent anti-MRSA (methicillin-resistant *Staphylococcus aureus*) activity. This novel structural modification potentially enhances bioactivity by inhibiting bacterial RNA polymerase, making it effective against various bacterial infections.

Another significant discovery includes <u>myxoquaterines</u>, a newly discovered family of natural products derived from an unusual combination of nonribosomal peptide synthetase (NRPS), polyketide synthase (PKS), and terpene biosynthesis. The myxoquaterines exhibit a DKxanthene-like core motif comprising a pyrrole and a methyloxazoline system linked by a double bond and followed by an olefinic side chain. This structural configuration implies a biosynthetic pathway related to that of DKxanthenes, though with significant evolutionary modifications. The stereochemistry includes multiple stereogenic centers, with NOE and H,H coupling constants suggesting a trans configuration for the methyloxazoline hydrogens and a half-chair conformation for the terpenoid ring in compound 4. Although myxoquaterine and DKxanthene pathways share structural and genetic elements, the former has undergone significant evolutionary changes that have resulted in their unique bioactivities. Myxoquaterines, particularly compound 4, exhibit significant bioactivity, which shows strong antiviral properties against HCoV-229E (IC50 340 ng/mL) and potent cytotoxicity against HCT-116 colon cancer cells (IC50 930 pg/mL) and KB3.1 cells (IC50 78 ng/mL).

Their genomic analysis is also supported by our comparative study revealing that all four *Pendulispora* organisms encode the highest numbers of BGC and NRPS-PKS pathways among the myxobacteria and all available organisms in the antiSMASH database **(Figure 1B, Table S2, Table S3)**. This indicates a substantial potential of their secondary metabolite at encoding and secretion levels, which warrants further exploration in future studies.

Overall, the identification of novel genes/proteins and the discovery of novel natural products in Pendulisporaceae, as mentioned in this article, highlights the immense potential of this new myxobacterial family for biotechnology and pharmaceutical applications. The unique BGCs and bioactivities of compounds like desferrioxamine PS, sorangicin P, myxoquaterines, and pendulipeptides underscore the value of investigating phylogenetically distant and underrepresented taxa. Further research into these novel natural products and their biosynthetic pathways could lead to the development of new antimicrobial and therapeutic agents.

## Supplementary Table

**Table S1) The distribution of BGC regions at Phylum level:** All BGC regions from >34000 organisms were extracted from the antiSMASH database and their median values, total number of organisms, and maximum/minimum number of BGC regions were calculated per phylum.

**Table S2) The distribution of BGC regions at Genus level:** All BGC regions from >34000 organisms were extracted from the antiSMASH database and their median/average values, total number of organisms, and maximum/minimum number of BGC regions were calculated per genus. The taxonomy of each genus is reported at domain, phylum, order, and family levels. Bar plots in column H represent organisms encoding >30 BGC regions at the median level.

**Table S3) The distribution of NRPS-PKS-hybrid regions at the Genus level:** The antiSMASH database was used to extract all BGC regions from >34000 organisms followed by the filtering of NRPS-PKS-hybrid regions. Their median/average values, total number of organisms, and maximum/minimum number of NRPS-PKS-hybrid regions were calculated per genus. The taxonomy of each genus is reported at domain, phylum, order, and family levels. Bar plots in column H represent organisms encoding >14 NRPS-PKS-hybrid regions at the median level.

## Ethics approval and consent to participate

Not applicable

## Availability of data and materials

Authors have used open-source tools in this analysis. All tool versions have been provided in the methodology.

## Authors’ contributions

Gaurav Sharma generated the idea. Gyanesh Shukla performed the analysis. Gaurav Sharma and Gyanesh Shukla edited and finalized the manuscript.

## Reference

1. Lewis, K. (2020). The Science of Antibiotic Discovery. Cell 181, 29–45. 10.1016/J.CELL.2020.02.056.

2. Hoskisson, P.A., and Seipke, R.F. (2020). Cryptic or silent? The known unknowns, unknown knowns, and unknown unknowns of secondary metabolism. MBio 11, 1–5. 10.1128/MBIO.02642-20/ASSET/DB6D3795-B353-4F01-B8D3-285C55C13534/ASSETS/GRAPHIC/MBIO.02642-20-F0001.JPEG.

3. Gould, K. (2016). Antibiotics: from prehistory to the present day. J. Antimicrob. Chemother. 71, 572–575. 10.1093/JAC/DKV484.

4. Shukla, R., Peoples, A.J., Ludwig, K.C., Maity, S., Derks, M.G.N., De Benedetti, S., Krueger, A.M., Vermeulen, B.J.A., Harbig, T., Lavore, F., et al. (2023). An antibiotic from an uncultured bacterium binds to an immutable target. Cell 186, 4059-4073.e27. 10.1016/J.CELL.2023.07.038.

5. Ziemert, N., Alanjary, M., and Weber, T. (2016). The evolution of genome mining in microbes – a review. Nat. Prod. Rep. 33, 988–1005. 10.1039/C6NP00025H.

6. Panda, A., Islam, S.T., and Sharma, G. (2022). Harmonizing Prokaryotic Nomenclature: Fixing the Fuss over Phylum Name Flipping. MBio. 10.1128/MBIO.00970-22.

7. Herrmann, J., Fayad, A.A., and Müller, R. (2017). Natural products from myxobacteria: novel metabolites and bioactivities. Nat. Prod. Rep. 34, 135–160. 10.1039/C6NP00106H.

8. Santos-Aberturas, J., and Vior, N.M. (2022). Beyond Soil-Dwelling Actinobacteria: Fantastic Antibiotics and Where to Find Them. Antibiot. 2022, Vol. 11, Page 195 11, 195. 10.3390/ANTIBIOTICS11020195.

9. Hoffmann, T., Krug, D., Bozkurt, N., Duddela, S., Jansen, R., Garcia, R., Gerth, K., Steinmetz, H., and Müller, R. (2018). Correlating chemical diversity with taxonomic distance for discovery of natural products in myxobacteria. Nat. Commun. 2018 91 9, 1–10. 10.1038/s41467-018-03184-1.

10. Blin, K., Shaw, S., Medema, M.H., and Weber, T. (2024). The antiSMASH database version 4: additional genomes and BGCs, new sequence-based searches and more. Nucleic Acids Res. 52, D586–D589. 10.1093/NAR/GKAD984.

11. Garcia, R., Popoff, A., Bader, C.D., Löhr, J., Walesch, S., Walt, C., Boldt, J., Bunk, B., Haeckl, F.P.J., Gunesch, A.P., et al. (2024). Discovery of the Pendulisporaceae: An extremotolerant myxobacterial family with distinct sporulation behavior and prolific specialized metabolism. Chem 0. 10.1016/j.chempr.2024.04.019.

12. Yamamoto, E., Muramatsu, H., and Nagai, K. (2014). Vulgatibacter incomptus gen. nov., sp. nov. and Labilithrix luteola gen. nov., sp. nov., two myxobacteria isolated from soil in Yakushima Island, and the description of Vulgatibacteraceae fam. nov., Labilitrichaceae fam. nov. and Anaeromyxobacteraceae fam. nov. Int. J. Syst. Evol. Microbiol. 64, 3360–3368. 10.1099/IJS.0.063198-0/CITE/REFWORKS.

13. Mohr, K.I., Garcia, R.O., Gerth, K., Irschik, H., and Müller, R. (2012). Sandaracinus amylolyticus gen. nov., sp. nov., a starch-degrading soil myxobacterium, and description of Sandaracinaceae fam. nov. Int. J. Syst. Evol. Microbiol. 62, 1191–1198. 10.1099/IJS.0.033696-0.

14. Sharma, G., Khatri, I., and Subramanian, S. (2016). Complete Genome of the Starch-Degrading Myxobacteria Sandaracinus amylolyticus DSM 53668T. Genome Biol. Evol. 8, 2520–2529. 10.1093/gbe/evw151.

15. Garcia, R., Gemperlein, K., and Müller, R. (2014). Minicystis rosea gen. nov., sp. nov., a polyunsaturated fatty acid-rich and steroid-producing soil myxobacterium. Int. J. Syst. Evol. Microbiol. 64, 3733–3742. 10.1099/IJS.0.068270-0.

16. Pal, S., Sharma, G., and Subramanian, S. (2021). Complete genome sequence and identification of polyunsaturated fatty acid biosynthesis genes of the myxobacterium Minicystis rosea DSM 24000T. BMC Genomics 22, 1–15. 10.1186/S12864-021-07955-X/FIGURES/5.

